# Width of Range Overlap in Congeneric Bird Species Pairs Along An Elevational Gradient In The Eastern Himalayas

**DOI:** 10.64898/2025.12.19.695388

**Authors:** Debjyoti Dutta, Rohan Pandit, Ramana Athreya

## Abstract

We investigated the distribution of range overlap between contiguous pairs of congeneric bird species along a 200-2800 m elevational gradient in the eastern Himalayas. Over a 3-year field survey between from 2012 to 2014 we recorded 15867 birds comprising 245 species in elevational intervals of 50 m. From this data set, we selected 107 congeneric pairs and fit a logistic curve to the abundance ratio versus elevation data to estimate the width of transition zone over which the species ratio changes from 90% to 10%. We found that the distribution of the transition widths range from < 50 m (about a quarter of the pairs) to more than 500 m. This lack of physical exclusion suggests that there is considerable niche separation between closely related species. For a smaller subset of species pairs with better quality data and model fit we investigated the correlation between the degree of range overlap (transition width) on the one hand, and phylogenetic distance and morphological trait difference on the other. We found a significant correlation with mass, beak width and wing size, but not with tarsus length, beak length and phylogenetic distance. This suggests that mass beak width and wing size may play an important role in niche differentiation in the area.

## 1. Introduction

The range of a species is defined as the geographical extent of a species occurrence and the area occupied by populations of a species (Gaston, 1991). Species ranges offer key insights into its ecological niche and variation in response to environmental change. The borders of a species range do not extend infinitely and tend to stabilise where environmental gradient is steepest (Case and Taper, 2000) often coinciding with locations where another species are more successful – either in terms of better adaptation to abiotic (environment) and/or biotic factors (competition, predation, pathogen pressures) (Holt and Keitt, 2005). Interspecific competition and gene flow have been theoretically shown to be the principal mechanisms involving biotic factors that act synergistically to set species range limits with competition reducing local densities of a species at it’s range edge and gene flow from central populations reducing the frequency of locally adapted alleles (Case and Taper, 2000).

The patterns of range boundaries arising as a result of such limiting mechanisms vary among species and have important ecological implications especially for congeneric species where range overlap is expected to induce competitive exclusion because of niche overlap. The extent of such overlaps indicate the degree of coexistence among congeners ultimately affecting their overall distribution. Range overlap patterns have also been shown to exhibit region specific variation in ecological properties of species as demonstrated in closely related species of birds whose body size differ more when their ranges overlap in warm as compared to cool climates (Bothwell, et al. 2015). Investigating species range overlap patterns therefore, provides crucial insight for predicting species persistence especially when climate change mediated range shift is a major concern.

Research on climate change induced range overlap involving closely related congenerics whose ranges do not currently overlap has predicted an overall future overlap for 6.4% of new world birds, mammals and amphibians with highest for birds (11.6%) followed by mammals (4.4%) and amphibians (3.6%) (Krosby et al. 2015). Future species overlap is expected to threaten persistence through increased competition and hybridisation (maladaptation for locally adapted individuals at range edge).

Muster et al (2002) illustrated possible patterns of range boundaries arising between species pairs arising as a result of various abiotic and biotic factors. These patterns may be ascribed to differing scenarios of relative fitness mediated by biotic and abiotic factors

Here we report an investigation into the overlap patterns in the ranges of 107 pairs of congeneric bird species along a steep elevational gradient spanning 200-2800 m in the eastern Himalayas. We estimated the width of the overlap zone of each pair. Anticipating that the overlap may be influenced by interspecific competition we also tested any relationship between the width of overlap and other parameters related to competition including phylogenetic distance and morphometric traits.

## 2. Methods

### 2.1 Study Location

The study was conducted along a 200-2800 m elevational transect in Eaglenest wildlife sanctuary in the eastern Himalayas of West Kameng District in Arunachal Pradesh, India (Athreya, 2006). Eaglenest is a small sanctuary of 218 km^2^ but has within it elevations ranging from 200-3250 m. The rainfall in this region ranges from about 1500 mm on the northern slopes to over 3000 mm on the southern slopes (Choudhury, 2003). The area was largely unexplored until the last two decades. Recent work has shown that the area is very rich in biodiversity and has been the focus of several research groups (Athreya, 2006; Srinivasan et al, 2018; Mungee & Athreya, 2021, Mungee et al. 2025). In particular, over 400 species of birds have been reported from the area.

The area between 500 and 2800 m is served by a motorable dirt track which facilitates access into the area. All bird records were obtain from this track which passes in the close proximity of primary forest.

### 2.2 Bird data

The bird data used in this study has been described elsewhere (Mungee & Athreya, 2021; Mungee et al, 2025; Maitra et al, 2022). Briefly, birds were recorded on road transects of 200 m length at 49 elevations between 200 m and 2800 m. The transects were separated by 50 m in elevation. Individual birds were identified to species and recorded by a single individual (Rohan Pandit). Each transect was sampled on 24 times during the summer months of May to July. This effort yielded 15867 records of 245 species.

The taxonomic identification of bird species initially followed Rasmussen and Anderton (2005). This paper follows the nomenclature and taxonomy Handbook of Birds of the World (HBW) and Birdlife International (http://www.birdlife.org/).

we identified 107 congeneric pairs of species for analysis from the 245 species in the data set.

### 2.3 Estimation Transition Width

We modeled the relative abundance of the two species using a logistic curve with elevation as the independent variable. The ratio of the abundance of the lower elevation species to that of all the individuals of the species pair (also termed the probablity of an individual being the lower species) was the dependent variable. The sigmoid function that we fit was p = 1/(1 + exp(β_0_ + β_1_E), where E is the elevation. We defined transition width as the elevational span over which the probability (of the lower elevation species) changed from 90% to 10%.

All analysis were done using scripts written in R (v4.3.2). The package glm in base R was used to model logistic regression.

### 2.4 Correlation between Overlap Width and Functional Traits

From the original list of 107 congeneric pairs we identified a subset of 52 species pairs for which the logistic fit was good, and both pairs had sufficient data (abundance > 10). We tested this restricted set for correlation between transition width on the one hand, and phylogenetic distance and morphometric traits on the other. The morphometric traits included body mass, beak length, tarsus length, beak width and wing length.

Phylogenetic distances for bird pairs were obtained from Schumm et al (2020) but there were 13 species in our dataset which were absent in Schumm et al’s (2020) dataset. These values were obtained from Mungee et al (2021) who had calculated them using the latest global phylogeny (Jetz et al, 2012). Morphometric trait values were obtained from Schumm et al (2020). We tested the relative value of the trait against transition width. The relative value of each trait was defined as the absolute difference of the values for the species pair, taken as a percentage of the maximum of the two.

## 3. Results

A representational sample of the elevational profiles of the ratio of abundances of congeneric species pairs are shown in Figures 1-3. In every plot, the x-axis is elevation while y-axis is the fraction (or probability) of an individual bird being of the lower elevation species in the congeneric pair. The plots also show the best fit logistic curve.

**Figure 1:**
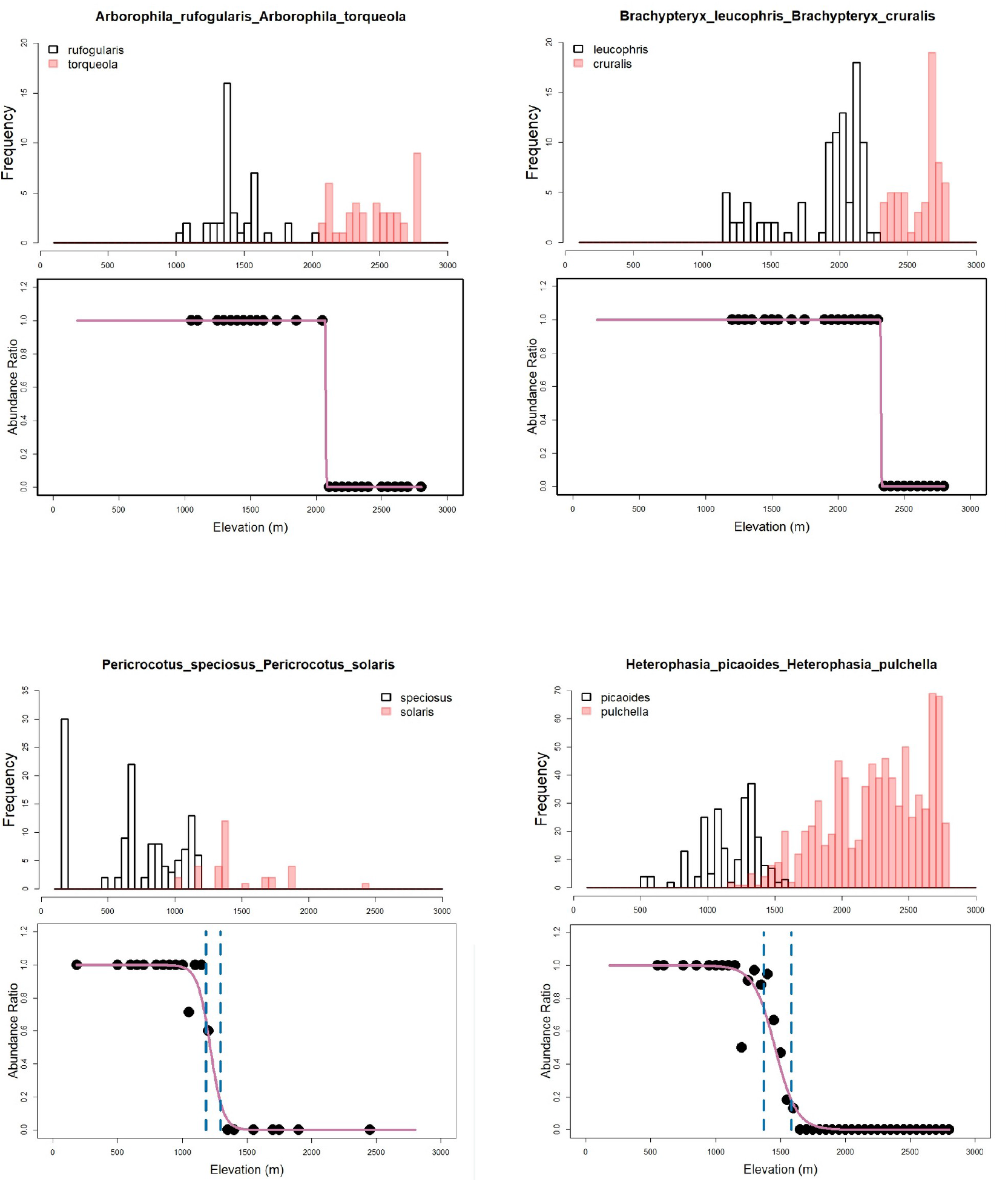
Logistic fit to abundance ratio of congeneric bird species.. The transition zone, wherein the abundance ratio falls from 90% to 10% is shown by vertical dashed line. These pairs are from the “good fit” category.

We identified 5 categories of range overlap based on the nature of the fit. They are listed in Table 1. The Figures mentioned earlier illustrate each of these types. The types are

**Table 1:**
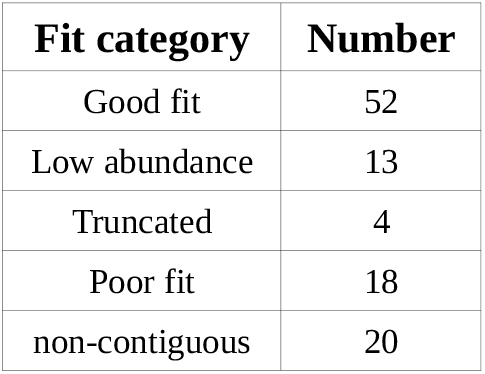
Category of logistic regression fits for abundance ratios of congeneric bird species pairs.

| Fit category | Number |
| --- | --- |
| Good fit | 52 |
| Low abundance | 13 |
| Truncated | 4 |
| Poor fit | 18 |
| non-contiguous | 20 |

We identified 5 categories of range overlap based on the nature of the fit. They are listed in Table 1. The Figures mentioned earlier illustrate each of these types. The types are

- “good” fits showed an overlap and the fit had a regular sigmoid shape (Figure 1)
- “low abundance” indicated very few individuals recorded (less than 10) and hence statistically not very reliable (Figure 2, bottom row)
- “truncated” fits were on edge of the sampled elevational range. They showed a downturn from the upper level but its proximity to the edge indicated that the estimated parameters may not be accurate (Figure 2, top row)
- “poor” fit had no discernible sigmoid shape to the curve (Figure 3, top row)
- “non-contiguous” were pairs in which there was no overlap and the distant between the closest records was more than 200 m (Figure 3, bottom row)

**Figure 2:**
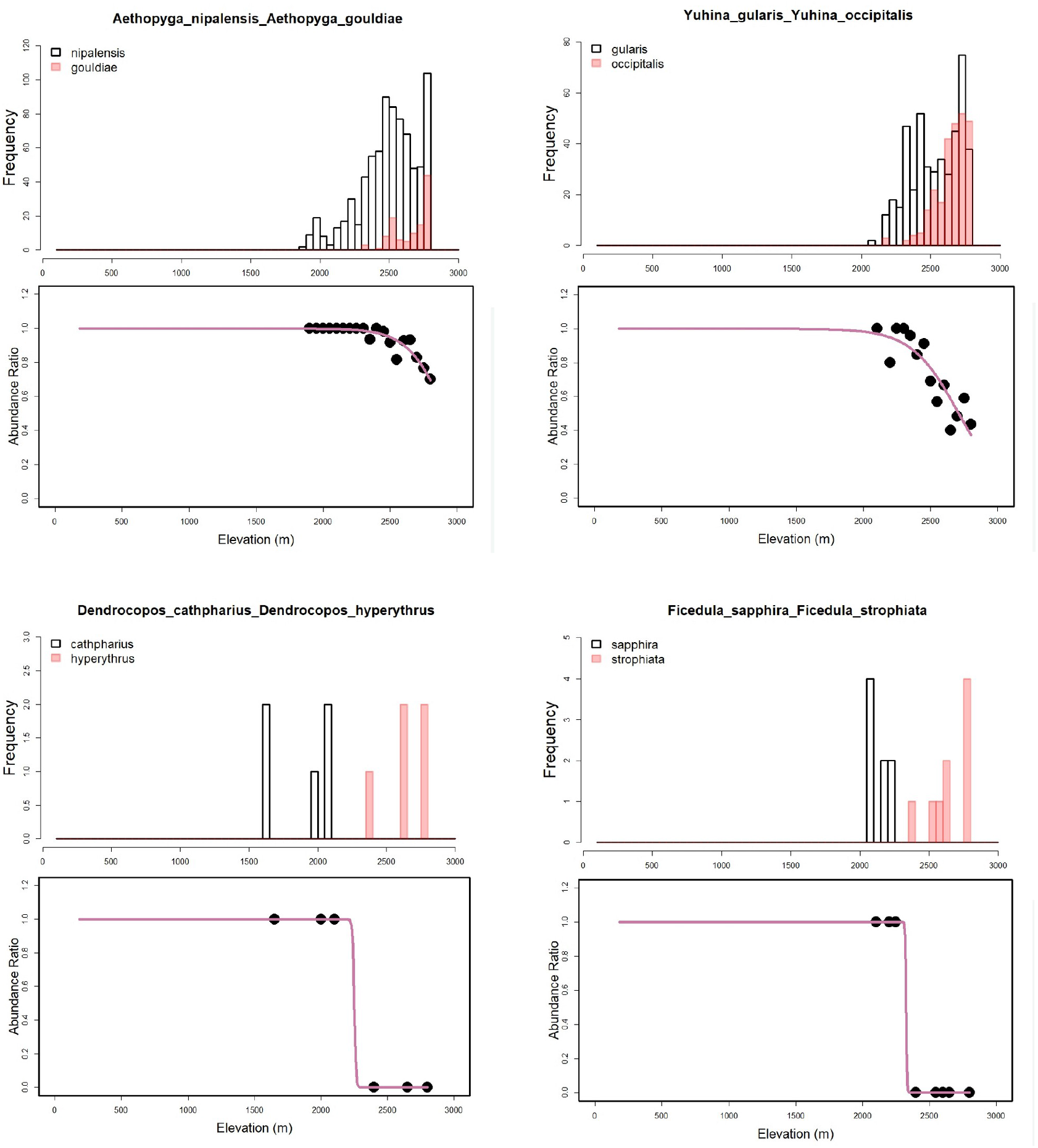
Logistic fit to abundance ratio of congeneric bird species. The upper plots are in the “truncated” category. The lower plots are “Too few” category.

**Figure 3:**
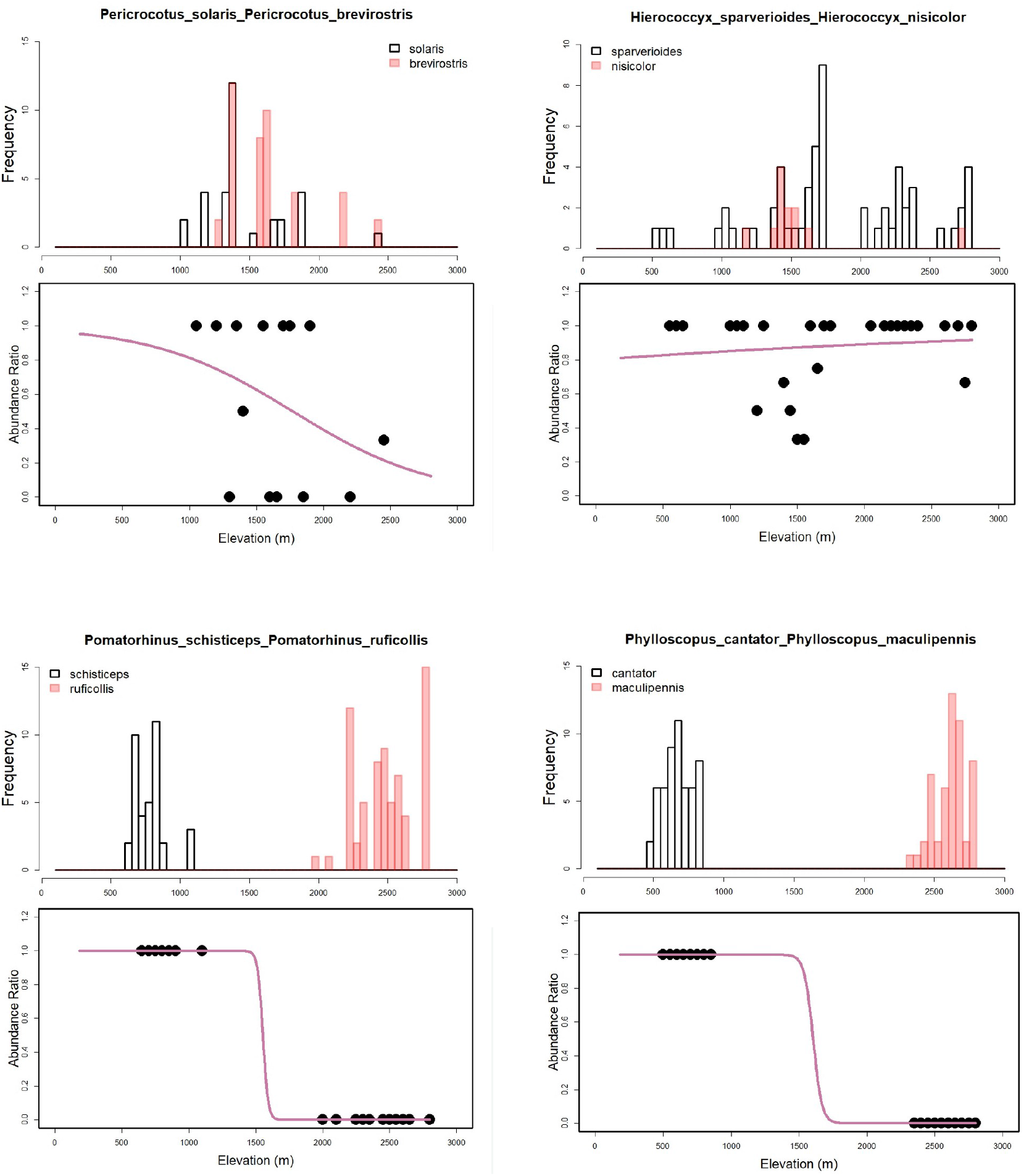
Logistic fit to abundance ratio of congeneric bird species. The upper plots are in the “poor fit” category. The lower plots are “non-contiguous” category.

While some of the “poor” fits are due to ecological factors – like broad ranges – henceforth, unless otherwise mentioned all discussion will be based on the species pairs with good logistic regression.

The distribution of transition widths is shown in Figure 4. There is no particular concentration of transition widths at small values in the “good” fit sample. Only a quarter of the species pairs have narrow transition widths of less than 100 m.

**Figure 4:**
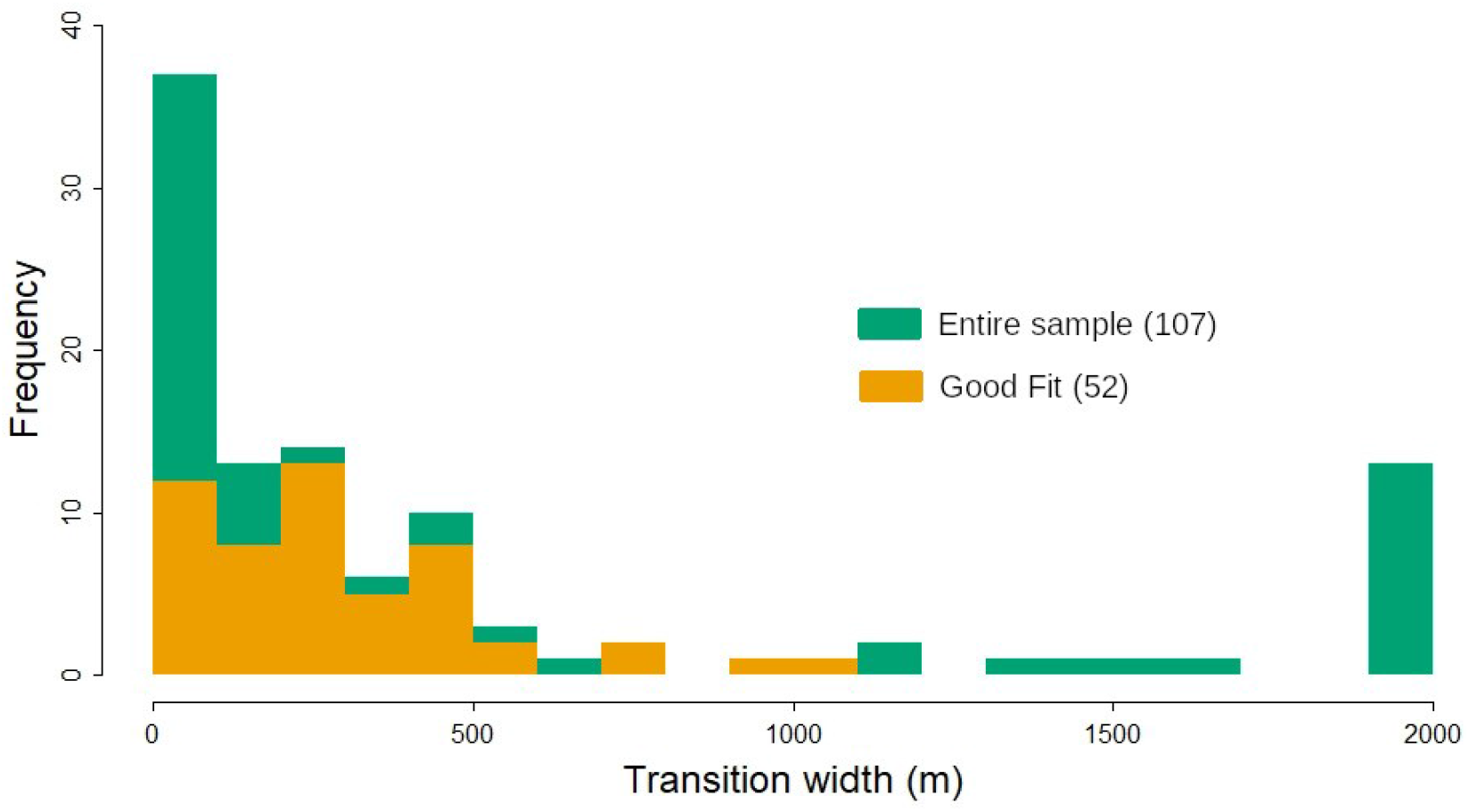
Histogram of transition width of 107 congeneric bird pairs. Widths greater than 2000 have been brought to 2000 for compactness. Overlapping orange bins represent transition width of 69 pairs selected for analysing relationship between morphometric traits and transition width.

Figure 5 shows the linear regression of transition width against morphometric traits.

**Figure 5:**
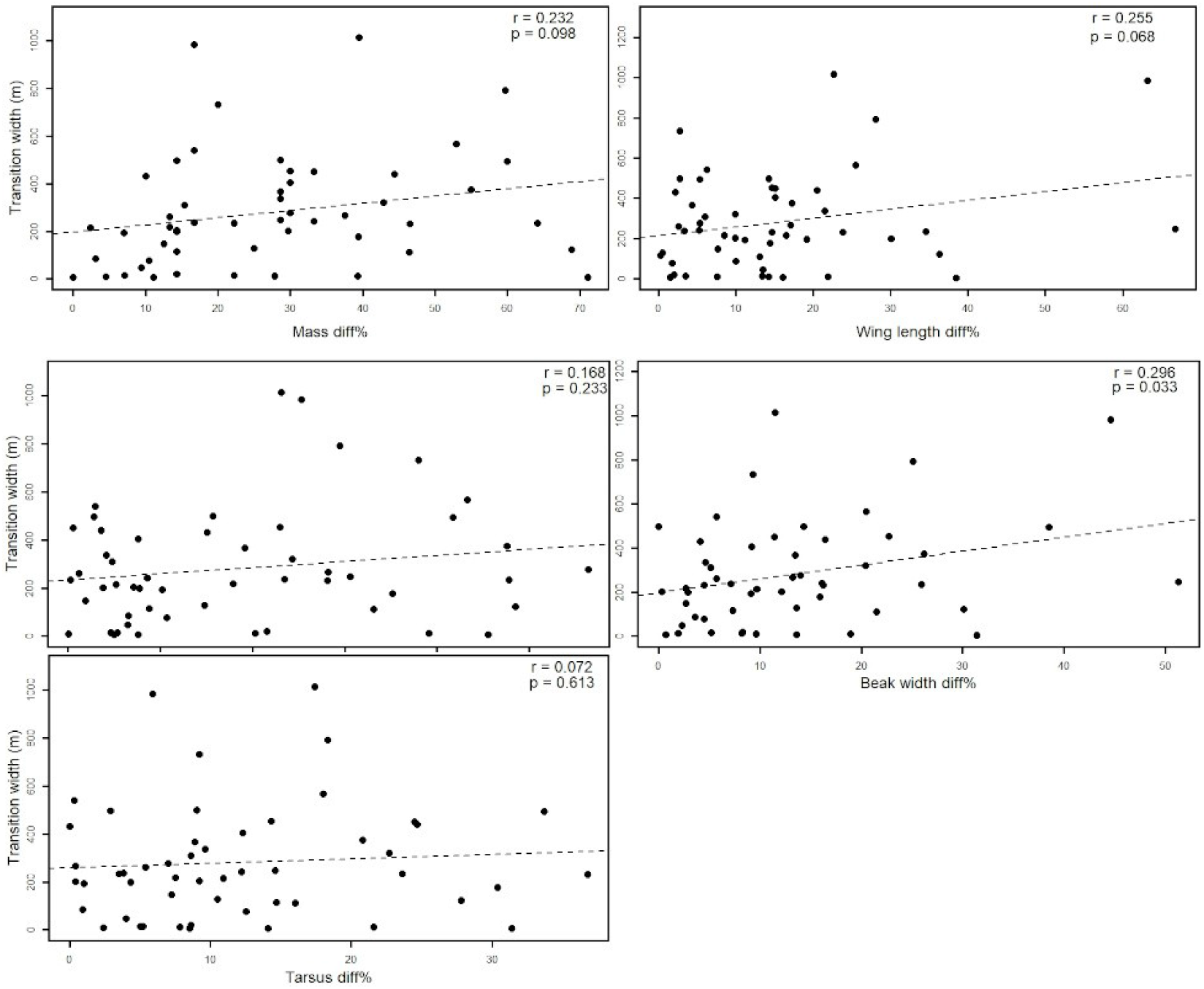
Linear regression of transition width and morphological traits.

The statistics of the regressions of transition width is listed in Table 2. Beak width shows a correlation at the 95% cofidence level. Mass and wing length show the same at a lower level of significance.

**Table 2:** Parameters of linear regression between transition width and other variables including phylogenetic distance and morphometric traits.Note that the correlation is with the percentage difference in trait between the species pair

| Predictor variable | Corr-coeff (r) | p-value |  |
| --- | --- | --- | --- |
| Phylogenetic Distance | 0.17 | 0.24 |  |
| Mass | 0.23 | 0.10 | * |
| Beak length | 0.17 | 0.23 |  |
| Tarsus length | 0.07 | 0.61 |  |
| Beak width | 0.30 | 0.03 | ** |
| Wing length | 0.26 | 0.07 | * |

The correlation means that greater the difference in the trait values, the wider the overlap in the species ranges.

Figure 6 shows the linear regression of transition width against phylogenetic distance. Three of the species pairs, which were supposedly congenerics, turned out to have very large phylogenetic distance. The plot on the right shows the regression without those 3 data points. There is no significant correlation between transition width and phylogenetic distance.

**Figure 6:**
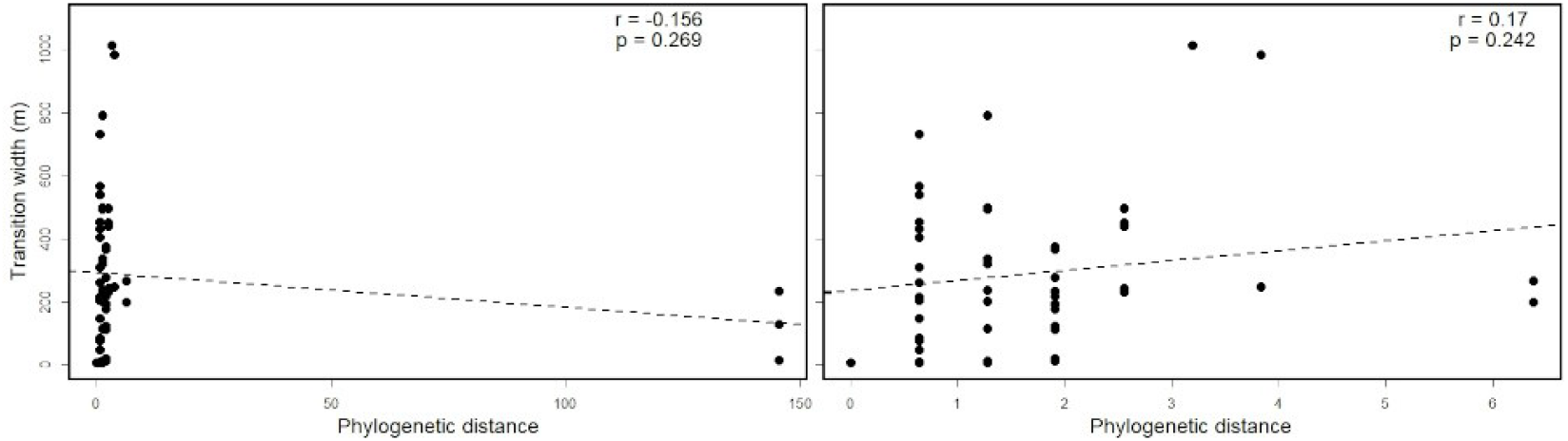
Linear regression of transition width and phylogenetic distance. Three supposedly congeneric species had very large phylogenetic distance (left). The analysis was redone without those 3 species pairs (right).

## 4. Discussion

Patterns of range overlap in elevationally distributed bird communities has been addressed in previous research as well. Elsen et al’s (2017) work investigating the distribution structure and range limits of birds found species range to be mostly constrained by temperature followed by ecotone and competition. Bird species whose ranges were limited by competition displayed similar patterns of elevational replacement of it’s congener but those limited by temperature and ecotone had overlapping ranges indicating species-wise variation in the relative importance of biotic and abiotic factors in generating range limits.

Srinivasan et al (2018) compared elevational distribution of bird communities between seasonal species-poor western himalayas and aseasonal species-rich eastern himalayas. They found that western species to be structured more by temperature and eastern species to be structured more by competition.

The major mechanism generating range limits among congeneric bird species of nightingle-thrushes distributed along low, mid and high elevation found competition to be the chief driver of range limit (Jankowski et al, 2010). The mid elevation species was found to react more aggressively to it’s high elevation counterpart’s song which can shrink the latter’s range more in the face of climate mediated range shifts.

Investigation of underlying factors affecting bird range depicted variation in causal factors with some finding territoriality instead of phylogenetic distance and morphometric trait as the main driving forces (Freeman et al. 2019; Freeman et al. 2022), some concluding physiological constraints establishing range limits (Altshuler, 2006) and some finding competition to be generating range limits in mountains (Cadena & Loiselle 2007). These investigations illustrate the variation in range overlap patterns among bird communities and shed’s light upon their probable causal mechanisms and it is this context which makes our work with birds relevant.

Our results suggest a species pair with a higher difference in flying ability also seem to be able to have a higher probability of co-existence. A higher ability to fly, indicated by larger wings, may indicate the ability to use a larger resource area. One can only speculate if a more vagile bird finds it easier to utilise resources in the territory of a less vagile bird.

On the other hand, mass and beak width are very important determinants of bird niches. A greater difference between the species pairs in these trait values would directly reduce competition between them. It is easier to interpret the correlation of these trait differences with the degree of range overlap.

The lack of a statistically significant relationship of range overlap and phylogenetic distance and the other two traits does not lend itself to a ready explanation. It could either mean that the traits are not very important in separating the niches or species, or more likely, niche separation is achieved by a complex combination of traits. That 3 traits showed a correlation is the surprise.

